# Serial Dependence in Position Perception Occurs at the Time of Perception

**DOI:** 10.1101/270272

**Authors:** Mauro Manassi, Alina Liberman, Anna Kosovicheva, Kathy Zhang, David Whitney

## Abstract

Observers perceive objects in the world as stable over space and time, even though the visual experience of those objects is often discontinuous and distorted due to masking, occlusion, camouflage, noise, etc. How are we able to easily and quickly achieve stable perception in spite of this constantly changing visual input? It was previously shown that observers experience serial dependence in the perception of features and objects, an effect that extends up to 15 seconds back in time. Here, we asked whether the visual system utilizes an object’s prior physical location to inform future position assignments in order to maximize location stability of an object over time. To test this, we presented subjects with small targets at random angular locations relative to central fixation in the peripheral visual field. Subjects reported the perceived location of the target on each trial by adjusting a cursor’s position to match its location. Subjects made consistent errors when reporting the perceived position of the target on the current trial, mislocalizing it toward the position of the target in the preceding two trials (Experiment 1). This pull in position perception occurred even when a response was not required on the previous trial (Experiment 2). In addition, we show that serial dependence in perceived position occurs immediately after stimulus presentation, and is a fast stabilization mechanism that does not require a delay (Experiment 3). This indicates that serial dependence occurs for position representations and facilitates the stable perception of objects in space. Taken together with previous work, our results show that serial dependence occurs at many stages of visual processing, from initial position assignment to object categorization.

## Introduction

Localization is one of the fundamental purposes of vision. Whether hunting for prey, looking for our wallet, or simply finding the computer mouse on our desk, correctly spotting an object’s location is essential to everyday life. Localization has been shown to depend on several factors, including motion (De Valois & De Valois, 1991; Ramachandran & Anstis, 1990; Whitney & Cavanagh, 2003, see also Whitney, 2002 for a review), spatial attention (Kerzel, 2000; Suzuki & Cavanagh, 1997), frames of reference (Bridgeman, Peery, & Anand, 1997), and eye movements (Cai, Pouget, Schlag-Rey, & Schlag, 1997; Ross, Morrone, & Burr, 1997; Ross, Morrone, Goldberg, & Burr, 2001). Perceived position can also be influenced by stimulus history. For example, adaptation to location (e.g., adaptation to information such as luminance distribution or textures that define object position) and adaptation to motion can shift the perceived location of a subsequent stimulus (Bressler & Whitney, 2006; McGraw, Whitaker, Skillen, & Chung, 2002; Nishida & Johnston, 1999; Snowden, 1998; Whitaker, McGraw, & Levi, 1997; Whitney & Cavanagh, 2003). These negative position aftereffects could reflect a mechanism that maximizes the visual system’s sensitivity to change (Gepshtein, Lesmes, & Albright, 2013).

Although sensitivity to change is clearly a useful function of vision, over-sensitivity to change may not be a universally desirable state; if sensitivity were too high, it might lead to a jittery or unstable perception of object position. Moreover, the world around us is generally static and autocorrelated: objects tend to remain in the same locations over time. The visual system may therefore balance the need to maximize sensitivity to change with the likelihood that the world is relatively stable, and such a mechanism could facilitate the stable perception of object location.

Recent work has hypothesized a novel mechanism for object stabilization, suggesting that perception occurs through continuity fields: spatiotemporally tuned operators within which similar features and objects are integrated (Fischer & Whitney, 2014). Continuity fields operate by inducing serial dependence in perception, making similar (but distinct) objects in time appear more similar than they actually are, and thus promoting the perception of object stability. For example, perceived orientation is systematically attracted toward previously seen orientations (Fischer & Whitney, 2014). Serial dependence has been shown to shape the perception of a variety of other objects and features beyond simply orientation (Fischer & Whitney, 2014), including faces (Liberman, Fischer, & Whitney 2014; Taubert, Alais, & Burr, 2016), attractiveness (Kondo, Takahashi, & Watanabe, 2012; Taubert, Van der Burg, & Alais, 2016; Xia, Leib, & Whitney, 2016), ambiguous objects (Tafazoli, Di Filippo, & Zoccolan, 2012; Wexler, Duyck, & Mamassian, 2015), motion (Alais, Leung, & Van der Burg, 2017), ensemble coding of orientation (Manassi, Liberman, Chaney, & Whitney, 2017), numerosity (Cicchini, Anobile, & Burr, 2014; Corbett, Fischer, & Whitney, 2011), and has also been shown to support stable object identity perception when an object moves behind an occluder (Liberman, Zhang, & Whitney 2016).

It remains unknown whether continuity fields only affect feature and object information or operate also on spatial information. Here, we tested whether continuity fields can generate serial dependence in the perceived position of objects. We flashed a target grating at random iso-eccentric locations and asked observers to report its position. We found systematic mislocalizations of the target, such that its reported location of the grating was biased: mislocalized toward previous target locations observed up to 10 seconds in the past.

## Methods

### Participants

Participants were affiliates of UC Berkeley and provided written informed consent before participation. All participants had normal or corrected-to-normal vision, and all except two were naïve to the purpose of the experiment. Twelve subjects (six female) participated in Experiment 1, ranging in age from 19 to 33 years (mean 25.5, s.d. 5). Sixteen subjects (seven female) participated in Experiment 2, ranging in age from 19 to 34 years (mean 25, s.d. 6). Four subjects from Experiment 1 participated in Experiment 2. Twelve subjects (four female) participated in Experiment 3A, ranging in age from 18 to 34 years (mean 23, s.d. 6), and eleven subjects (five female) participated in Experiment 3B, ranging in age from 19 to 33 years (mean 22, s.d. 4). All experimental procedures were approved by the UC Berkeley Institutional Review Board and were in accordance with the Declaration of Helsinki.

### Stimuli & Procedure

Experiments were conducted in a darkened experimental booth. Subjects viewed stimuli on a CRT monitor (1024 x 768, 100 Hz, Dell Trinitron) at a distance of 56 cm. All experiments were programmed in Matlab (The MathWorks, Natick, MA) with Psychophysics Toolbox (Brainard, 1997).

Subjects were instructed to continuously fixate a central black dot (0.21° in diameter) on a gray background (21 cd/m^2^) during the experiment. Observers were presented with a series of target stimuli (linear grating inside a circular aperture) at random locations on an iso-eccentric circle (10° eccentricity; Figure 1A). Each target stimulus consisted of a vertically-oriented static sine wave grating (4 cycles per degree, Michelson contrast 30%) embedded in a circular mask with a 3° diameter (hard aperture). In order to increase difficulty in detecting the position of the Gabor, pink noise (1/f) was added to the grating. The position of the grating was randomized across 48 possible positions, in rotation steps of 7.5° (Figure 1B). On each trial, the target was shown for 80 ms, followed by a pink noise mask (150 ms) that filled the entire screen. The purpose of the mask was to minimize any possible negative aftereffect from the grating. Next, a black dot (0.5° diameter) appeared in a random location on the iso-eccentric circle, also at 10° eccentricity. Subjects were asked to adjust the black dot’s position with the mouse to match the position of the grating. The dot was constrained to only move clockwise and counterclockwise along the invisible iso-eccentric circle. The next trial began after an inter-trial interval (ITI) of 2500 ms. A long ITI was used in order to maintain a similar trial duration to Fischer & Whitney (2014). The fixation dot was present for the entire duration of the trial, including the ITI. Each observer completed 450 trials in total.

Experiment 2 was the same as Experiment 1, except that in 33% of the trials, observers were not asked to perform the position adjustment task. Rather, observers were presented with an inter-stimulus interval (ISI) of 1250 ms, in which only the fixation dot was shown. The purpose of this experiment was to remove the response requirement and investigate whether motor response biases might play a role in any potential serial position effect. Each observer completed 1380 trials in total.

Experiments 3A and 3B were the same as Experiment 1, except that no mask was presented after the grating stimulus. In Experiments 3A and 3B, the gratings had a Michelson contrast of 30% and 4%, respectively.

**Figure 1:**
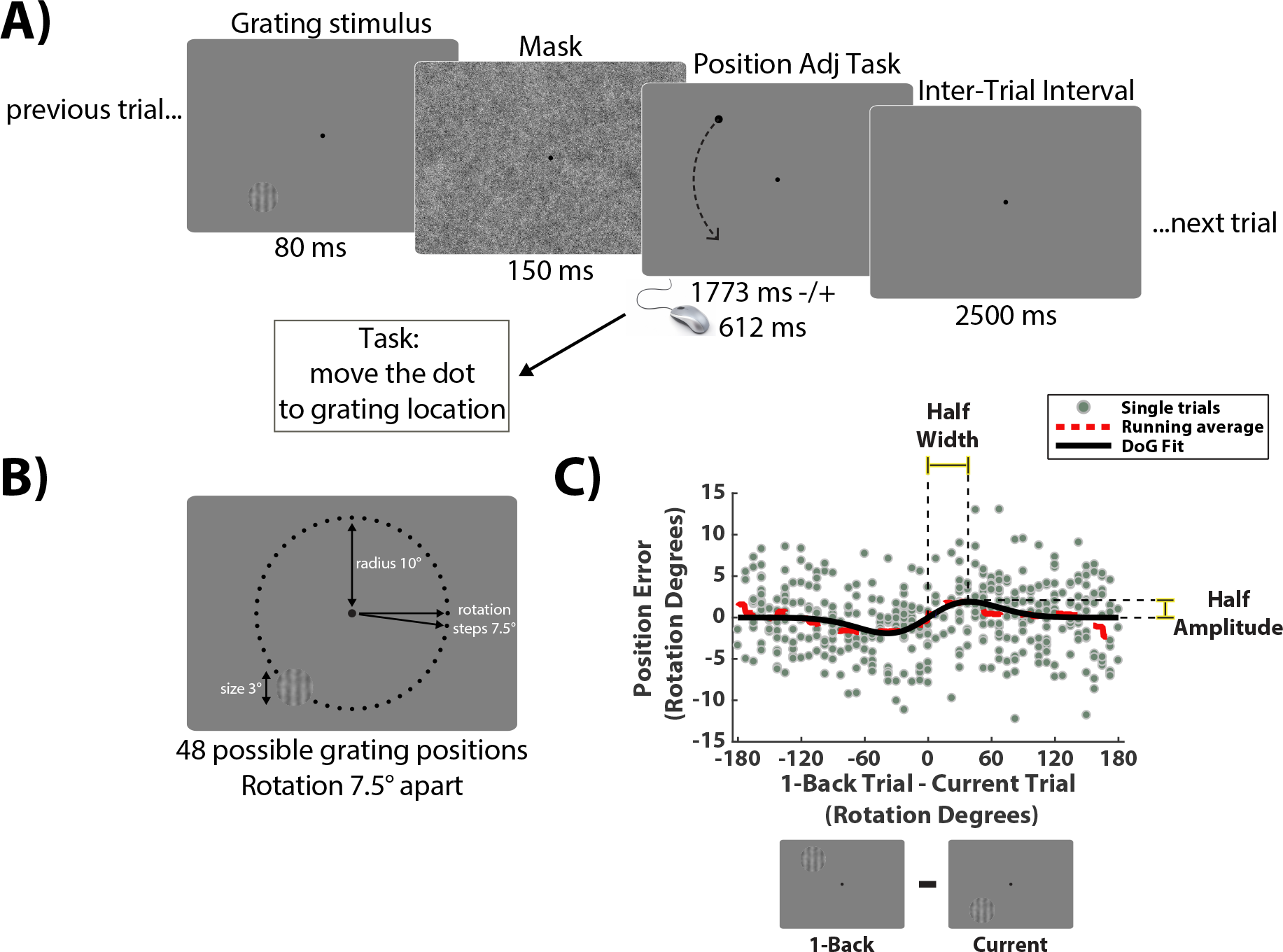
Trial sequence in Experiment 1. A) Observers were instructed to fixate on a central dot during the experiment. A target grating was presented at a random location on an invisible iso-eccentric circle for 80 ms. After a noise mask of 150 ms (to reduce afterimages), subjects were asked to report the perceived location of the target by adjusting a dot’s position to match the location of the grating. Trials were separated by a 2500 ms delay. B) The grating (3° diameter) was presented at random angular locations relative to central fixation, at an eccentricity of 10°. There were 48 possible positions, in rotation steps of 7.5°. C) Example data from a representative subject (1 trial back). Each data point shows performance on one trial. The x-axis represents the difference between the previous position 1 trial back and the current position. The y-axis represents the error in the adjustment task (difference between dot position in the adjustment task and grating position on current trial). The average error (red line) shows more negative (counterclockwise) response errors for a negative relative position and more positive (clockwise) errors for a positive relative position. In order to quantify the magnitude of serial dependence, we fit a derivative-of-Gaussian (DoG) to the data (black line) measuring the half-amplitude peak for each observer.

### Data analysis

For each subject’s data, trials were considered lapses and excluded if the error exceeded 3 s.d. from the grand mean error in perceived position, calculated for each individual subject, or if the response time (RT) was longer than 10 s (less than 5% of data excluded on average). Response error for each trial was computed as the angular difference between the subject’s response, given by the position of the adjustment dot and the actual position of the grating (Figure 1C; y-axes; reported target position minus actual target position). Position was always determined by the rotation angle from the fixation point at the center of the screen. Negative and positive values indicate that the subject’s response was more counterclockwise or clockwise relative to the actual grating, respectively. Response error was then compared to the difference between the current and previous grating position (Figure 1C, x-axes; target position on previous trial minus target position on current trial). Negative and positive values indicate that the previous grating was in a more counterclockwise or clockwise position compared to the current grating, respectively.

In order to quantify the strength of serial dependence, we fit a simplified Gaussian derivative (DoG) to each subject’s data (Figure 1C) of the form:

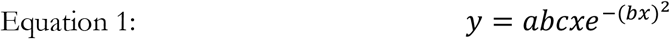

where 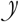 is response error on each trial, x is the relative orientation of the previous trial, a is half the peak-to-trough amplitude of the derivative-of-Gaussian, b scales the width of the Gaussian derivative, and c is a constant 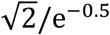, which scales the curve to make the *a* parameter equal to the peak amplitude. We fit the Gaussian derivative using constrained nonlinear minimization of the residual sum of squares. As a measure of serial dependence, we report half the peak-to-trough amplitude (parameter *a*; Figure 2A, 3, 4) and half the width (parameter *b*; Figure 2B) of the best fitting derivative-of-Gaussian. A positive value for the *a* parameter indicates a perceptual bias toward the position of the previous grating. A negative value for the *a* parameter indicates a perceptual bias away from the position of the previous grating. A value of zero for the *a* parameter indicates no bias.

Kosovicheva & Whitney (2017) recently showed that that individual subjects have idiosyncratic biases in reported position unrelated to serial dependence, such as perceptual distortions at different points around the isoeccentric stimulus circle. For this reason, we conducted an additional control analysis to remove such potential unrelated biases before fitting the Gaussian derivative function described in Equation 1. To do this, we fit a polynomial function (10 degrees) to each observer’s error distribution (reported target position minus actual target position) as a function of target location on the circle. Systematic motor error, for example, might manifest as a bias to consistently report a target presented at the 12 o’clock position as being at the 2 o’clock position. To regress out such biases, we subtracted the subject’s reported target positions from the discretized polynomial fit. This subtraction left us with residual errors that did not include the idiosyncratic biases unrelated to serial dependence. We then plotted these residual errors as a function of the difference between current and previous target location (x-axes in Figure 1C) and fit the Gaussian derivative to the data. Importantly, the addition of this control analysis—removing systematic biases unrelated to serial effects—had no significant impact on the serial dependence results. It did not generate or increase the measured serial dependence.

## Results

### Experiment 1: Serial dependence in perceived position

Ten subjects out of twelve displayed a positive DoG half-amplitude, indicating that perceived position on a given trial was significantly pulled in the direction of grating position on the preceding (i.e., 1-back) trial (t(11) = 2.94, p = 0.013, Figure 2A). Even when comparing subjects’ errors with the difference in position two trials back, subjects showed significantly positive DoG half-amplitudes (t(11) = 3.82, p = < 0.01, Figure 2A), meaning that even grating positions presented two trials back biased perceived position on a given trial. For three and four trials back, DoG half-amplitudes were not significantly different from zero (3-back: t(11) = 1.51, p = 0.15; 4-back: t(11) = 0.47, p = 0.64; Figure 2A). Average response time across subjects was 1773 ± 612 ms. The perceived position of the grating was therefore strongly attracted toward previous grating positions seen 5 seconds (1 trial-back, Figure 2A) or 10 seconds ago (2 trials-back, Figure 2A).

We also analyzed the width of the DoG fit in order to address whether the temporal tuning of serial dependence (1-2 trials back) determines its spatial tuning (i.e., the width in the DoG fit). There were no statistically significant differences in width between 1, 2, 3 and 4 trials back as determined by one-way ANOVA (*F*(3,44) = 0.09, *p* = .96). In addition, there was no correlation between half amplitudes and half width (Figure 2B; r = −0.16, p = 0.26).

**Figure 2:**
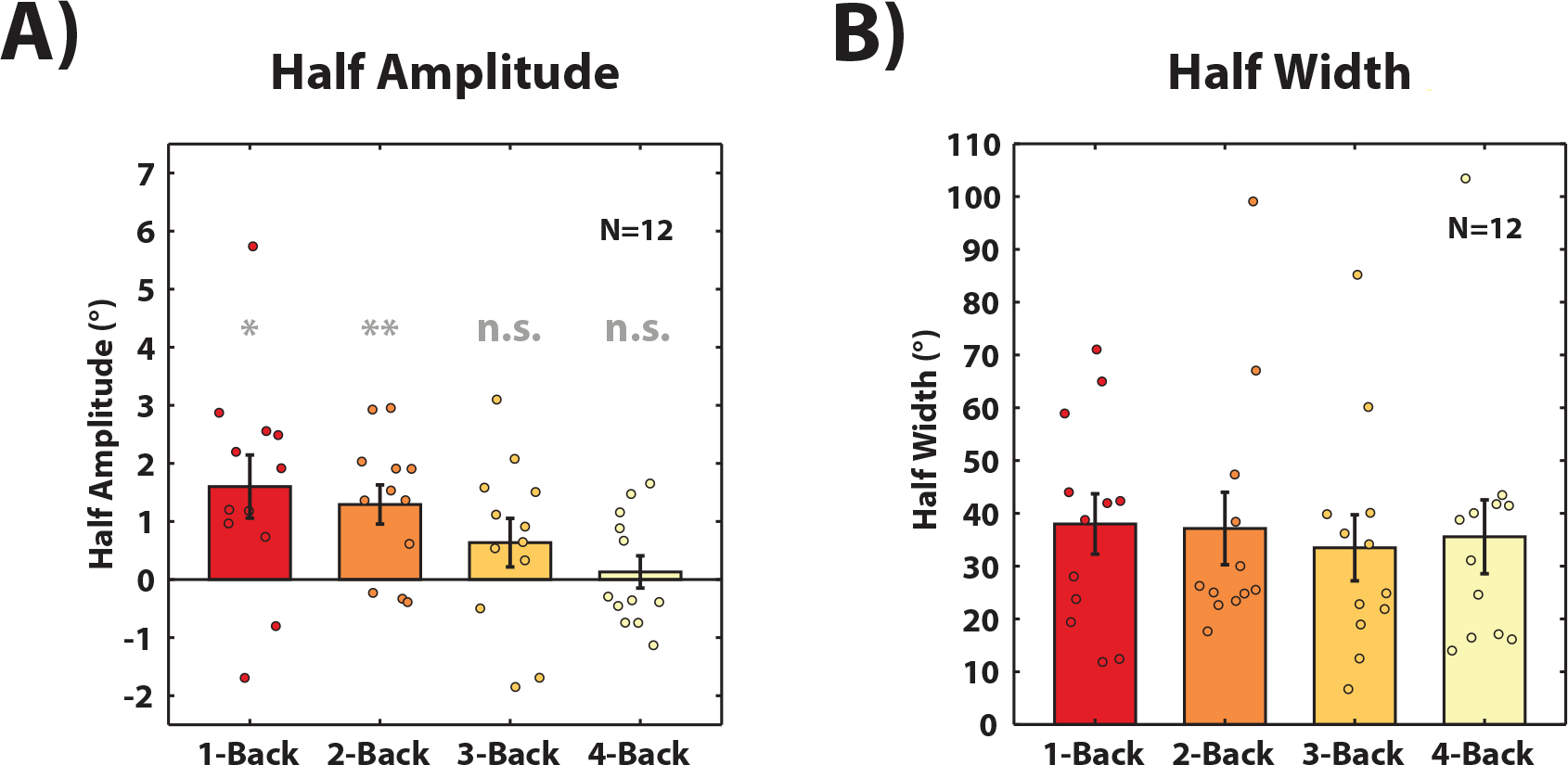
Experiment 1. Average half amplitudes (A) and widths (B) of Derivative of Gaussian fit across twelve subjects for 1, 2, 3 and 4 trials back. Each colored dot represents the half amplitude or width for a single subject (horizontal jitter added for visibility). Bars indicate the average and error bars indicate ± one standard error. The asterisks represent a significant difference from zero.

### Experiment 2: Serial dependence in position is not due to previous motor response

In Experiment 1, we found that subjects made consistent errors when reporting the perceived location of the target grating on the current trial, mislocalizing it toward the location presented on the previous two trials.

In Experiment 2, we tested whether this position bias may be due to the previous motor response. In this experiment, on 33% of the trials, the position adjustment task was replaced with an ISI of 1250 ms. Importantly, position adjustment task trials (67% of trials) and no-task trials (33% of trials) were presented in a random fashion, and observers did not know whether they would have been asked to respond or not to the target in a given trial. If serial dependence observed in Experiment 1 was due to the previous motor response, then we should expect no evidence of serial dependence in the position adjustment task if it was preceded by a trial containing an ISI in place of the position adjustment task. Each subject completed 1380 trials divided into 12 blocks.

In both sequences (Figure 3), subjects displayed on average positive DoG half-amplitudes (previous response: t(15) = 3.94, p < 0.001; no previous response: t(15) = 2.58, p = 0.02), meaning that serial dependence in perceived position did not require a previous motor response (Figure 3, red and purple bars). We also found increased serial dependence when the previous trial required a response (Figure 3, red bars) than when it did not (Figure 3, purple bars; t(15) = 3.87, p < 0.001). This increase in effect observed in the response conditions was likely due to the fact that subjects were presented with an additional dot at the same grating position, thus reinforcing the serial dependence effect.

As an alternative analysis, we could have measured serial dependence from the previous response by computing the x-axis (Figure 1C) as “adjusted dot position in previous trial (previous response rather than previous stimulus) minus grating position on current trial”. However, in this specific analysis, motor response biases (Shaffer, 1978; Wing & Kristofferson, 1973), oblique effects (Appelle, 1972; Cicchini, Mikellidou, & Burr, 2017; Mikellidou, Cicchini, Thompson, & Burr, 2015), or other localization biases (e.g., Kosovicheva & Whitney, 2017) can be consistent and therefore correlate across trials. Any of those consistent biases can create artifacts which resemble serial dependence, but are in fact unrelated. Because of these methodological issues, previous studies have measured serial dependence between current and previous stimuli (Cicchini et al., 2017; Fischer & Whitney, 2014; Liberman et al., 2014; Manassi et al., 2017), rather than comparing previous and current response.

**Figure 3:**
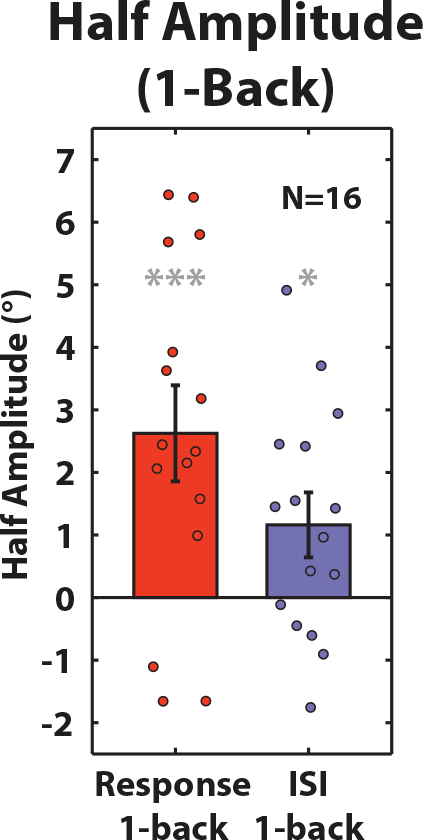
Experiment 2. Average half amplitudes of DoG when previous trial required a response (previous response, red bar) and when previous trial was interleaved with an ISI of 1250 ms (no response, purple bar).

### Experiment 3A and 3B: Fast serial dependence in position perception

Recently, it was proposed that serial dependence in position perception is a phenomenon of mnemonic rather than perceptual processes (Bliss, Sun, & D’Esposito, 2017). With a very similar paradigm^1^, the authors varied the delay between stimulus presentation (a very high contrast black dot) and response (dot position adjustment) and found that positive serial dependence occurred for delays between the target dot and the adjustment dot that ranged from 1 to 10 seconds. Interestingly, when there was no delay between the target and adjustment dot (and no mask), there was a negative aftereffect instead of positive serial dependence (see Figure 2B in Bliss et al., 2017). It is known that positive serial dependence and adaptation-induced negative aftereffects can be additive (Alais et al., 2017; Cicchini et al., 2017; Fischer & Whitney, 2014; Taubert, Alais, et al., 2016), and the negative aftereffect Bliss et al. (2017) report could be the result of adaptation to the high contrast, salient target dot, as previous authors have reported (Hess, Dakin, & Badcock, 1994; Whitaker et al., 1997). To reduce adaptation and negative aftereffects, Experiment 1 used a mask following the stimulus. In Experiment 3A and 3B, we removed the mask and varied the contrast of the target in order to test whether serial dependence occurs when there is zero delay between stimulus presentation and response.

Experiment 3A was the same as Experiment 1, except that no noise mask was presented after the target grating presentation. The noise mask was removed in order to keep the interval between target and response stimulus at zero. Michelson contrast of the grating was kept at 30% as in Experiments 1 and 2 (high contrast experiment). Each observer completed 450 trials in total.

Subjects showed no evidence of serial dependence or negative aftereffect. The DoG half-amplitudes were not significantly different from zero for any trial back (1-back: t(11) = 1.30, p = 0.21; 2-back: t(11) = 2.01, p = 0.07; 3-back: t(11) = 0.67, p = 0.51; 4-back: t(11) = −1.40, p = 0.18; Figure 4A). Taken together, these results are in accordance with previous results (Bliss et al., 2017), showing no serial dependence for a 0 delay between stimulus and response (Experiment 3A), and serial dependence for longer delays (Experiments 1-2). However, this does not mean that the delay, per se, is the critical factor.

The critical factor that modulates serial dependence in perceived position may be the effective contrast of the target stimulus. Previous evidence showed that the perceived contrast of a grating can be weakened by a subsequent (backward) mask (Breitmeyer, Rudd, & Dunn, 1981; Breitmeyer, Hoar, Randall, & Conte, 1984; Breitmeyer & Ogmen, 2000; Kolers, 1962; Raab, 1963) and, as a consequence, its negative aftereffect is weakened (Gibson & Radner, 1937; Keck, Palella, & Pantle, 1976; Stecher, Sigel, & Lange, 1973). Accordingly, a grating with a backward mask showed serial dependence (Experiments 1 and 2), whereas a grating without mask showed no evidence of serial dependence (Experiment 3A). Hence, if the apparent reduction in target contrast (and not the delay) determines the strength of serial dependence, we hypothesized that serial dependence should arise with a lower contrast grating. In Experiment 3B we tested this hypothesis.

In Experiment 3B, we reduced the Michelson contrast of the grating to 4% (low contrast experiment). Our aim was two-fold. First, by reducing contrast, any adaptation induced negative aftereffect (Hess et al., 1994; McGraw et al., 2002; Whitaker et al., 1997) would be reduced. Second, subjects are forced to pay more attention to the target grating. As serial dependence strongly depends on attention (Fischer & Whitney, 2014), a stronger positive bias is expected.

In accordance with our hypothesis, subjects showed evidence of serial dependence, even with zero delay between the target and response stimulus. This held for 1 and 2 trials back (1-back: t(10) = 5.18, p < 0.001; 2-back: t(10) = 9.36, p < 0.001; 3-back: t(10) = 2.07, p = 0.06; 4-back: t(10) = 1.15, p = 0.27; Figure 4B). Hence, the critical factor that determines the strength of serial dependence in localization judgments is the luminance contrast (or uncertainty more generally), not the delay. Serial dependence in position is a fast mechanism, biasing position perception immediately after stimulus presentation. Although serial dependence may still be modulated by short-term working memory (Bliss et al., 2017; Fritsche, Mostert, & de Lange, 2017), it cannot be considered a purely delay-dependent or working memory based process detached from its perceptual component (see also Cicchini et al., 2017).

**Figure 4:**
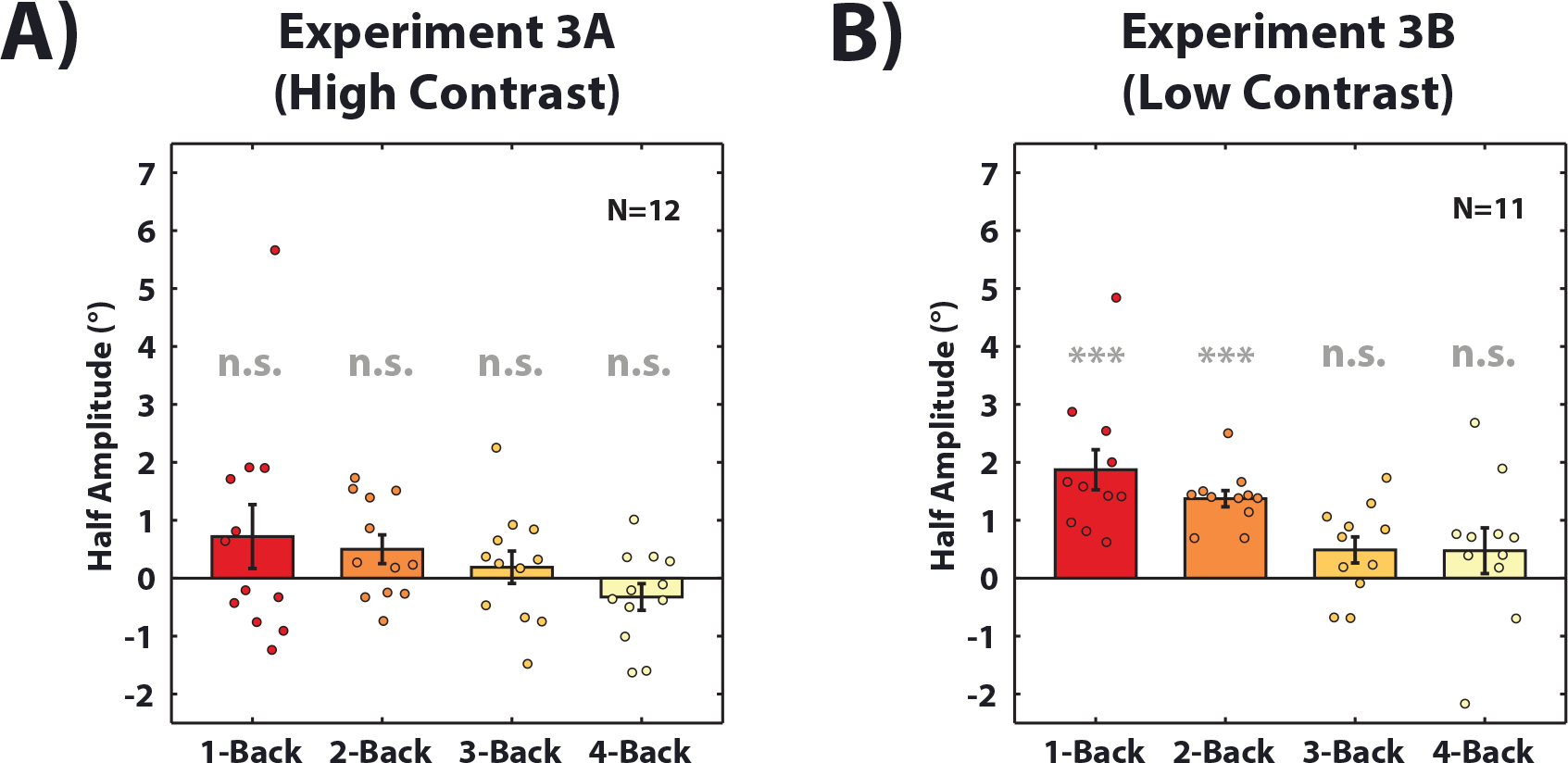
Experiment 3A and 3B. Average half amplitudes of Derivative of Gaussian fit for 1, 2, 3 and 4 trials back. Grating contrast was 30% in Experiment 3A and 4% in Experiment 3B. Each colored dot represents the half amplitude for a single subject. Bars indicate the average and error bars indicate ± one standard error. The asterisks represent a significant difference from zero.

## Discussion

The world around us appears stable despite changes in noise and lighting, discontinuities such as eye blinks, and changes in gaze and head position. Previous studies have proposed that, in order to facilitate perceptual stability, perception occurs through continuity fields: spatio-temporal regions over which object are attracted toward recently seen objects and features (Fischer & Whitney, 2014; Liberman et al., 2014). On a neural level, continuity fields may advantageously reduce cortical processing and re-processing from moment-to-moment, by recycling representations of previously perceived features and objects. On a perceptual level, continuity fields help observers maintain a continuous and stable representation of the world.

Recent papers have also suggested that there is serial dependence across species. Papadimitriou, Ferdoash, & Snyder (2015) found that when monkeys made saccades to remembered positions, the saccades were biased toward previous saccade targets. Along the same lines, Tafazoli et al. (2012) found that when rats were trained to learn the appearance of a default object, they perceived two new subsequent objects as similar to the previous default one. Our results are consistent with and extend these prior studies, showing that serial dependence occurs in human spatial vision, even in tasks as fundamental as perceptual localization.

To summarize our findings across the experiments reported here, we found (A) a positive aftereffect in position: position perception was pulled by object positions encountered five or ten seconds ago (Experiment 1; Figure 2). (B) This kind of serial dependence is not due to the subject’s motor response in the previous trial; when we eliminated the motor response in the 1-back trial, we still found a significant serial dependence effect on position in the current trial (Experiment 2; Figure 3). (C) Serial dependence in position perception occurs immediately, independent on the delay between stimulus and response (Experiment 3A-3B; Figure 4). Taken together, our results suggest that serial dependence can occur in position representations, providing a mechanism through which continuity fields can promote the appearance of position consistency from moment to moment.

Our results may involve serial dependence at the level of spatial memory representations (Papadimitriou et al., 2015; Rahnev, Koizumi, McCurdy, D’Esposito, & Lau, 2015). In fact, there may be a strong connection between serial dependence in perception and in memory (Kiyonaga, Scimeca, Bliss, & Whitney, 2017; see also Makovski & Jiang, 2008; Papadimitriou et al., 2015). Serial dependence has a clear perceptual component (Cicchini et al., 2017; Fischer & Whitney, 2014), but higher level factors like decision (de Lange & Fritsche, 2017; Fritsche et al., 2017) and memory (Bliss et al., 2017) may still play a role in determining its strength. It is also possible that there is serial dependence at multiple levels of representation, including at basic perceptual levels (Cicchini et al., 2017; Fischer & Whitney, 2014), perceptual decisions (Fritsche et al., 2017), and also in memory representations (Papadimitriou et al., 2015; Zhang, Liberman, & Whitney, 2016). How these multiple levels interact remains an exciting area of investigation.

Prior work has shown that previous trials can affect subsequent trials in detecting target position (Maljkovic & Nakayama, 1996) through priming. However, this priming effect manifests primarily as a reduction in reaction times: reaction times decreased when the target position was repeated on consecutive trials, and increased when the target fell on a position previously occupied by a distractor. Conversely, our results showed that previous positions biased the reported location of the target. While it is still debated whether serial effects are due to a change in perception or decision, continuity fields underlie all these effects for the same purpose: to promote stable localization in the complex environments we experience everyday (Fischer & Whitney, 2014; Manassi et al., 2017).

Previous work on perceptual localization has shown that adaptation to motion (McGraw et al., 2002; Nishida & Johnston, 1999; Snowden, 1998; Whitaker et al., 1997; Whitney & Cavanagh, 2003; Whitney, 2005) and luminance- or texture-defined stimuli (Hess et al., 1994; McGraw et al., 2002; Nishida & Johnston, 1999; Snowden, 1998; Whitaker et al., 1997) can lead to a *repulsion* effect, biasing perceived visual position away from previously seen adaptors. Our results reveal an *attraction* toward previous positions. The visual system implements adaptation and repulsion mechanisms in order to maximize sensitivity to change or differences. Our results indicate that there is a complementary attraction mechanism that facilitates perceptual stability of position. These two mechanisms could be opposite sides of the same coin: both facilitate constancy and stability, but do so differently. Open questions remain as to how adaptation (and repulsion effects more generally) interact with serial dependence, and some work has begun to investigate this (Fischer & Whitney, 2014; Taubert, Alais, et al., 2016). For example, there may be different time courses for adaptation and serial dependence, or a different dependence on noise, inter-stimulus intervals, or storage, among other factors. Future research should investigate the interplay between these two opposing mechanisms and how the balance between these processes facilitates our perception of stability.

In conclusion, our results provide evidence that serial dependence occurs for object localization. Together with previous work, this suggests that continuity fields determine visual perception at several stages of visual processing. Serial dependence has been shown to occur for low level features, such as position assignment and orientation perception (Fischer & Whitney, 2014; Fritsche et al., 2017; Liberman et al., 2016), as well as for high level features such as face perception (Liberman et al., 2014; Taubert, Alais, et al., 2016) and attractiveness (Kondo et al., 2012; Taubert, Van der Burg, et al., 2016; Xia et al., 2016). The existence of continuity fields at several stages of visual processing, along with their spatial and temporal tuning, suggests that the underlying neural mechanism(s) likely involves feedback, but is not likely to be a single stage process or a unitary decision effect.

## Acknowledgments

This work was supported in part by the Swiss National Science Foundation fellowship P2ELP3_158876 (M.M.) and NSF graduate research fellowships (NSF-GRFP) to A.K. and A.L This work was originally presented at Vision Science Society Annual Meeting in 2014. We would like to thank Daniel Bliss for useful discussions.

## Footnotes

1 Their work was independently submitted after our initial submission to Psychonomic Bulletin and Review.

